# Differential gene expression supports a resource-intensive, defensive role for colony production in the bloom-forming haptophyte, *Phaeocystis globosa*

**DOI:** 10.1101/461756

**Authors:** Margaret Mars Brisbina, Satoshi Mitaraia

## Abstract

*Phaeocystis globosa* forms dense, monospecific blooms in temperate, northern waters. Blooms are usually dominated by the colonial morphotype—non-flagellated cells embedded in a secreted mucilaginous mass. Colonial *Phaeocystis* blooms significantly affect food-web structure and function and negatively impact fisheries and aquaculture, but factors initiating colony production remain enigmatic. Destructive *P. globosa* blooms have been reported in tropical and subtropical regions more recently and warm-water blooms could become more common with continued climate change and coastal eutrophication. We therefore assessed genetic pathways associated with colony production by investigating differential gene expression between colonial and solitary cells in a warm-water *Phaeocystis globosa* strain. Our results illustrate a transcriptional shift in colonial cells with most of the differentially expressed genes downregulated, supporting a reallocation of resources associated with colony production. Dimethylsulfide and acrylate production and pathogen interaction pathways were upregulated in colonial cells, suggesting a defensive role for colony production. We identify several protein kinase signaling pathways that may influence the transition between morphotypes, providing targets for future research into factors triggering colony production. This study provides novel insights into genetic mechanisms involved in *Phaeocystis* colony formation and provides new evidence supporting a defensive role for *Phaeocystis* colonies.

## INTRODUCTION

*Phaeocystis* is a cosmopolitan bloom-forming haptophyte genus encompassing 6 species (Andersen et al. 2015; Schoemann et al 2005). Most *Phaeocystis* species (*P. globosa, P. antarctica, P. pouchetii*, and *P. jahnii*) exhibit a polymorphic life-cycle, alternating between colonial and free-living morphotypes. *Phaeocystis* blooms are usually dominated by the colonial morphotype and are typically very dense, produce large biomasses, and impact food-web structure and function (Schoemann et al. 2005). *Phaeocystis* is a major contributor to dimethylsulfoniopropionate (DMSP) and dimethylsulfide (DMS) production globally (Liss et al. 1994) and regional peaks in DMS production are closely correlated with colonial *Phaeocystis* blooms (Van Duyl et al. 1998). DMS produced in the surface ocean is aerosolized and its oxidation products promote cloud formation, increase albedo, and affect global climate (Charlson et al. 1987). In algal cells, DMSP and its cleavage products, DMS and acrylate, contribute to osmotic balance, neutralize reactive oxygen species, and deter grazing (Noordkamp et al. 2000; Sunda et al. 2002). Despite the ecological importance of colony formation in *Phaeocystis*, triggers for transition to the colonial morphotype remain enigmatic, and the functional role of colony formation in the *Phaeocystis* life-cycle is not clearly delineated (Peperzak & Gabler-Schwarz 2012).

Myriad factors have been studied in regard to their roles in instigating colony formation in *Phaeocystis* species, including nutrient and light availability (Bender et al. 2018; Cariou et al. 1994; Wang et al. 2011), temperature (Verity & Medlin 2003), mechanical stress (Cariou et al. 1994), grazing cues (Long et al. 2007; Tang 2003; Wang et al. 2015), and viral infection (Brussaard et al. 2005; Brussaard et al. 2007). However, these studies used different *Phaeocystis* species, strains, and morphotypes with a range of experimental conditions, which yielded variable and sometimes contradictory results. Nonetheless, several lines of evidence suggest that colony formation serves a defensive role. First, while viruses can cause 30-100% cell lysis in solitary *Phaeocystis*, viruses rarely infect colonial cells, which lyse primarily due to nutrient limitation (Brussaard et al. 2005; Brussaard et al. 2007). Second, ciliates and other microzooplankton that graze solitary *Phaeocystis* are unable to graze on colonies (Tang et al. 2001) and chemical cues from these grazers induce colony formation and promote increased colony size (Long et al. 2007; Tang 2003). Third, acrylate, which is produced with DMS when DMSP is cleaved, accumulates within colonies and may further deter macro- and micro-grazers and heterotrophic bacteria (Hamm et al. 2000; Noordkamp et al. 2000). However, while cellular growth rate increases in colonial cells relative to solitary cells if colonies are induced in nutrient rich conditions, it decreases when colonies are induced under nutrient limiting conditions (Wang et al. 2015). Thus, colony formation can defend against pathogens and grazers, but it is costly (Wang et al. 2015), suggesting that colony formation is likely a complex response to interacting biotic and abiotic factors (Long et al. 2007).

Colony formation may also play a fundamental role in *Phaeocystis* reproduction. *Phaeocystis* has one of the most complex and polymorphic life cycles among phytoplankton genera, and despite extensive study, it remains largely unresolved in most species. Studies have implicated at least 6 different life stages and up to 15 functional components to the life-cycle (Gaebler-Schwarz et al. 2010). In *P. globosa*, four morphotypes are believed to exist: diploid colonial cells devoid of scales and flagella, diploid scale-free flagellates arising from mechanically disrupted colonies, and two types of small, scaled, haploid flagellates—those that produce vesicles containing star-shaped alpha-chitin filaments and those that do not (Rousseau et al. 2007). Haploid flagellates may fuse (syngamy) to produce diploid colony-forming cells, which in turn undergo meiosis and produce haploid flagellates (Rousseau et al. 2013). Haploid flagellates are often observed swarming inside colonies, suggesting that colonial bloom formation may contribute to successful sexual reproduction in *Phaeocystis* (Peperzak et al. 2000; Rousseau et al. 2013). However, neither syngamy nor meiosis have been directly observed in *Phaeocystis* spp. (Peperzak & Gabler-Schwarz 2012), even though both events have been documented in several other haptophyte genera (Houdan et al. 2003). If colonial *Phaeocystis* blooms are necessary for sexual reproduction, it would further justify the resource costs associated with colony formation.

Historically, colonial *Phaeocystis* blooms have been restricted to cold, high-latitude waters—*P. globosa* blooms in the English Channel and North Sea, *P. pouchetii* in the North Atlantic and Arctic, and *P. antarctica* in the Southern Ocean (reviewed in Schoemann et al. 2005). In the last two decades, however, blooms have increasingly been reported in tropical and subtropical regions, including the subtropical N. Atlantic (Long et al. 2007) and the subtropical and tropical South China Sea (Chen et al. 2002; Doan-Nhu et al. 2010; Liu et al. 2015). Decaying colonial biomass sinks and produces anoxic conditions, making *Phaeocystis* blooms detrimental to benthic fisheries and aquaculture (Desroy & Denis 2008; Peperzak & Poelman 2008; Spilmont et al. 2009). In warmer waters, *Phaeocystis globosa* blooms have been especially catastrophic to local aquaculture (Chen et al. 2002; Doan-Nhu et al. 2010), possibly because the hemolytic activity of *P. globosa* liposaccharides increases with temperature (Peng et al. 2005). Global climate change and increasing nutrient pollution in coastal regions may mean harmful *Phaeocystis* blooms will continue to increase in range and frequency. Given the ecological impact of colonial *Phaeocystis* blooms and their complex and enigmatic initiating triggers, particularly in warm waters, it is imperative to better understand the regulation of colony formation.

Transcriptional approaches have become an exceptionally useful tool to illuminate physiological responses to environmental cues and genes associated with specific life-stages in algae and other protists (Caron et al. 2017). In this study, we investigated genetic regulation of colony formation by analyzing gene expression in colonial and flagellated morphotypes of a warm-water *Phaeocystis globosa* strain. Since *Phaeocystis* is an important marine producer of DMSP and DMS—and because these molecules may be associated with colonial defense, we queried our dataset for algal genes involved in DMSP production (*DSYB*, Curson et al. 2018) and its cleavage to DMS and acrylate (*Alma1*, Alcolombri et al. 2015). *DSYB* and *Alma1* are the only algal genes that encode proteins experimentally proven to catalyze DMSP, DMS, and acrylate production, but neither has been identified in *P. globosa* previously. Overall, our results demonstrate a dramatic transcriptional shift in colonial *P. globosa*, with the vast majority of differentially expressed genes downregulated in colonial cells. Such a strong transcriptional shift supports an allocation of resources toward colony formation and away from other cellular processes such as translation, cell growth, and cell division. Genes associated with DMSP production (*DSYB*-like) were not differentially expressed, but an *Alma* family-like gene was upregulated in colonies, suggesting colonies may produce more DMS and acrylate than do solitary cells. Results further indicate that pathogen interaction affects colony formation, further supporting a defensive role for colonies, and implicating several cell-signaling pathways as important to colony formation. This study provides new insights into the functional role of *Phaeocystis* colonies and physiological processes associated with colony formation. These insights will guide future investigations into triggers that initiate colony formation in harmful *Phaeocystis* blooms.

## MATERIALS AND METHODS

### Culture strain and maintenance

Starter cultures of *Phaeocystis globosa* CCMP1528, a warm-water, colony forming *P. globosa* strain (Wang et al. 2011), were purchased from the National Center for Marine Algae and Microbiota (NCMA, Maine, U.S.A.) in January 2016. CCMP1528 did not initially form colonies in our culture conditions, but it is known that *P. globosa* sometimes stops forming colonies in culture (Janse et al. 1996). We maintained non-colonial cultures in replicate 150 ml Erlenmeyer flasks with 100 ml L1-Si media in ambient light and temperature on a gently rotating twist mixer (TM-300, AS ONE, Osaka, Japan, speed setting 1). Culture media was prepared by enriching 34 ppt autoclaved artificial seawater prepared from milliQ water and sea salts (Marine Art SF-1, AS ONE, Osaka, Japan) with the NCMA L1 media kit (-Si) and filtering through sterile 0.22 μm pore-size filters. Cultures were diluted biweekly with freshly prepared media. When one culture replicate began producing colonies in January 2017, experimental culture conditions were promptly initiated.

### Experimental culture conditions

We prepared four biological replicates each of colony-forming and non-colonial *P. globosa* CCMP1528 by inoculating 45 ml of sterile L1 media with 1 ml stock culture in 50-ml Erlenmeyer flasks. Replicates were placed on a gently rotating twist mixer in a plant growth chamber with cool white fluorescent lamps (CLE-305, TOMY, Tokyo, Japan) set to 22 °C with light level 4 and a 12:12 day:night ratio. A HOBO temperature and light logger (Onset, MA, U.S.A.) was kept in the growth chamber during the experiment. The daytime temperature was 21 ° C with about 1900-2000 Lux (~30 μmol m^−2^ s^−1^) light intensity, and the nighttime temperature was 22 °C. Positions of replicates were rotated daily to prevent position in the chamber from systematically affecting replicates. On days 1, 3, and 4, chlorophyll fluorescence was measured at the middle of the light period by transferring 200 μl aliquots to a black 96-well plate (ThermoFisher, MA, U.S.A.) and recording fluorescence (excitation: 440 nm, emission: 685 nm) with a Tecan Ultra Evolution microplate reader (Tecan, Mannedorf, Switzerland). Exponential growth phase was determined by comparing measured fluorescence to a growth curve for *Phaeocystis globosa* CCMP 1528 grown under identical conditions. On day 4 of the first round, 1 ml of each replicate was transferred to 45 ml of sterile L1 media and the experimental setup was repeated, which allowed for adaptation to experimental culture conditions. Algal cells were harvested for RNA extraction on day 4 of the second experimental culture round, when replicates were in middle to late exponential growth phase (Fig. S1).

Prior to RNA extraction, each culture replicate was imaged with light microscopy (Olympus CKX53, MA, U.S.A.) to ensure that colony-forming replicates were indeed producing colonies and that non-colonial replicates were not (Fig. S2). Flagellates present in the colonial and non-colonial replicates were actively swimming, suggesting that flagellates in this study were scaled haploid flagellates, rather than scale-free diploid flagellates originating from disrupted colonies, but neither flow-cytometry nor electron microscopy were performed (Rousseau et al. 2007). Colony-forming culture replicates were filtered through polytetrafluoroethylene (PTFE) filters (10-μm pore size) (Millipore, NH, U.S.A.) under gentle vacuum. Colonies were visible by eye on the filter surface and swimming flagellates were observed in the flow-through when viewed with light microscopy. The non-colonial culture replicates were first filtered through 50-μm nylon mesh to remove culture debris, and then filtered through 1.0 μm pore-size PTFE filters. No flagellates were visible when the filtrates were viewed with light microscopy. Filters were immediately flash frozen in liquid nitrogen and stored at −80 °C until RNA extraction.

### RNA extraction, library preparation and sequencing

Total RNA was extracted from filters by following the manufacturer’s protocols for the MoBio PowerWater RNA extraction kit (Qiagen, MD, U.S.A.), including the optional initial heating step. Following extraction, we assessed RNA quality and concentration. RNA extracts were diluted so that 10 ng of RNA were used for each sample with the SMART-seq v4 Ultra Low Input RNA Kit (Clonetech/Takara, CA, U.S.A.) along with 2 μl of a 1:10,000 dilution of External RNA Controls Consortium (ERCC) spike-in mix 1 (Ambion, CA, U.S.A.), an internal quality control. The SMART-seq kit employs poly-A priming to target eukaryotic mRNA and to reduce the amount of ribosomal and bacterial RNA present in sequencing libraries. The quality and concentration of the resulting cDNA was assessed before continuing with the manufacturer’s protocols for the Nextera XT DNA Library Prep Kit (Illumina, CA, U.S.A.). Finally, we checked cDNA fragment size before submitting libraries to the Okinawa Institute of Science and Technology DNA Sequencing Section for paired-end 150×150 bp sequencing across 8 lanes of an Illumina Hiseq4000 flow-cell.

### Bioinformatic processing and quality control

Sequencing reads were processed with Trimmomatic software to remove adapter sequences and to filter low-quality sequences (Bolger et al. 2014). Read quality was checked with FastQC before and after trimming to ensure that adapters were removed (Andrews 2010). Remaining reads were mapped to the ERCC reference sequences (Cronin et al. 2004) and mapped reads were counted with RSEM software (Li & Collin 2011). Counts were further analyzed in the R statistical environment (R Core Team 2013). Reads mapping to the ERCC reference sequences were then removed from each sample with SAMtools (Li et al. 2009) and BEDTools (Quinlan & Hall 2010).

### Transcriptome assembly, assessment, and functional annotation

The Marine Microbial Eukaryote Transcriptome Sequencing Project (MMETSP, Keeling et al. 2014) assembled a transcriptome for *Phaeocystis sp.* CCMP2710, which groups with the *Phaeocystis globosa* species complex in phylogenetic analyses (Fig. S3). Only 25% of our reads, however, mapped to this reference transcriptome. We therefore assembled a de novo transcriptome for *Phaeocystis globosa* CCMP1528 to serve as a reference for read mapping in this study. We used Trinity software for transcriptome assembly (Grabherr et al. 2013) and dereplicated the transcriptome by removing reads with 95% similarity using CD-HIT-EST (Fu et al. 2012). Bacterial contamination was removed by performing a blastn query against the NCBI nucleotide database (downloaded March 2018, ncbi-blast v2.6.0+, Camacho et al. 2009) and parsing results to identify and remove bacterial contigs. The final assembly was assessed for completeness with Benchmarking Universal Single-Copy Orthologs (BUSCO v3, Simao et al. 2015) and results were compared with those for the MMETSP *Phaeocystis sp.* CCMP2710 transcriptome.

We annotated the CCMP1528 transcriptome using two different databases, Pfam (Finn et al. 2010) and KEGG (Kanehisa et al. 2016). Pfam annotation was performed with the dammit software (Scott 2018), which wraps Transdecoder to translate transcriptome contigs to the longest possible amino acid sequence (Haas et al. 2013), and HMMER to assign protein homologs to sequences (Eddy 2011). After discarding annotations with e-values greater than 1E-5, the annotation with the lowest e-value was selected for each contig. Gene Ontology (GO) terms were assigned to Pfam annotations using the Gene Ontology Consortium’s Pfam2GO mapping (geneontology.org/external2go/pfam2go, version 07/14/2018, Mitchell et al. 2015). KEGG annotation was performed with the GhostKOALA tool and translated amino acid sequences (kegg.jp/ghostkoala, 05/21/2018, Kanehisa et al. 2016). Annotated K numbers were then used to assign KEGG pathways by accessing the KEGG API (kegg.jp/kegg/rest/keggapi.html, July 2018).

### Differential gene expression analysis

Quality filtered sequences from each sample were mapped to the assembled *P. globosa* CCMP1528 transcriptome and counted with RSEM software. Counts for each sample were imported into the R statistical environment, where differential gene expression between colonial and solitary culture replicates was tested with the DESeq function in the Bioconductor package DESeq2 (Love et al. 2014). Genes that were differentially expressed were considered statistically significant if the False Discovery Rate (FDR) adjusted p-value (padj) was less than 0.05.

### Gene set enrichment testing

We identified GO terms enriched among significantly upregulated and downregulated genes by applying a hypergeometric test in the R package GOstats (Falcon & Gentleman 2007). GOstats accommodates user-defined GO annotations, which are necessary when studying non-model organisms like *Phaeocystis*. Likewise, a hypergeometric test for significant enrichment of KEGG pathways was applied using the enricher function from the R package ClusterProfiler (Yu et al. 2012). Because of the lower annotation rate, KEGG pathway enrichment was further investigated by additionally applying linear model analysis with the kegga function in the R package edgeR (Robinson et al. 2010). GO terms and KEGG pathways were considered significantly enriched when the statistical test returned a p-value less than 0.05.

### Genes associated with DMSP and DMS production

Because *Phaeocystis* is a profusive producer of DMSP and DMS, we specifically queried our dataset for recently discovered algal genes involved in DMSP production (DSYB) and its cleavage to DMS and acrylate (Alma family genes). We performed blastp queries with curated DSYB protein sequences (provided by Curson et al. 2018) and *E. huxleyi* and *Symbiodinium* Alma family protein sequences downloaded from UniProt (July 2018) against *Phaeocystis globosa* CCMP1528 amino acid sequences. Expression levels of putative *Phaeocystis globosa DSYB* and *Alma* family genes were then checked in colonial and solitary culture replicates.

## RESULTS

### Bioinformatic processing and quality control

Sequencing for this project produced over 1.9 billion read pairs with 159-383 million read pairs per sample. The reads for each sample were deposited in the Sequence Read Archive (SRA) with accession numbers SRR7811979-SRR7811986. Following quality filtering with Trimmomatic, 1.7 billion read pairs remained, with 140-341 million read pairs per sample (Table S1). After mapping reads from each sample to ERCC reference sequences, we plotted the Log_2_ FPKM for each sequence against the Log_2_ of its concentration in the standard mix. A simple linear regression was fitted for each sample and R^2^ values ranged from 0.93-0.937 for each sample (Fig. S4). The strong correlation between observed FPKM and initial concentration for ERCC sequences indicates that minimal bias was introduced during PCR amplification, library preparation, and sequencing.

### Transcriptome assembly, assessment, and functional annotation

The final assembly of the *Phaeocystis globosa* CCMP1528 transcriptome included 69,528 contigs and a total of 43.9 Mbp (available for download from https://doi.org/10.5281/zenodo.1476491). The CCMP1528 transcriptome was about 3 times larger than the MMETSP CCMP2710 transcriptome, but the minimum, maximum, and mean contig lengths were about the same for both (Table S2). When Transrate was used to align the two transcriptomes, only 18% of CCMP1528 contigs aligned to the CCMP2710 transcriptome, but 55% of the CCMP2710 contigs aligned to the CCMP1528 transcriptome. BUSCO software was utilized to assess completeness of the *P. globosa* CCMP1528 transcriptome. It included more complete eukaryote and protist BUSCOs than the MMETSP *Phaeocystis sp.* CCMP2710 transcriptome (Fig.1). Together, these results demonstrate that the transcriptome generated in this study is more complete than the MMETSP transcriptome and is a better reference for this study. The results also indicate that CCMP2710 and CCMP1528 are more genetically distant than was expected based upon ribosomal RNA gene sequences (Fig. S3).

**Figure 1.**
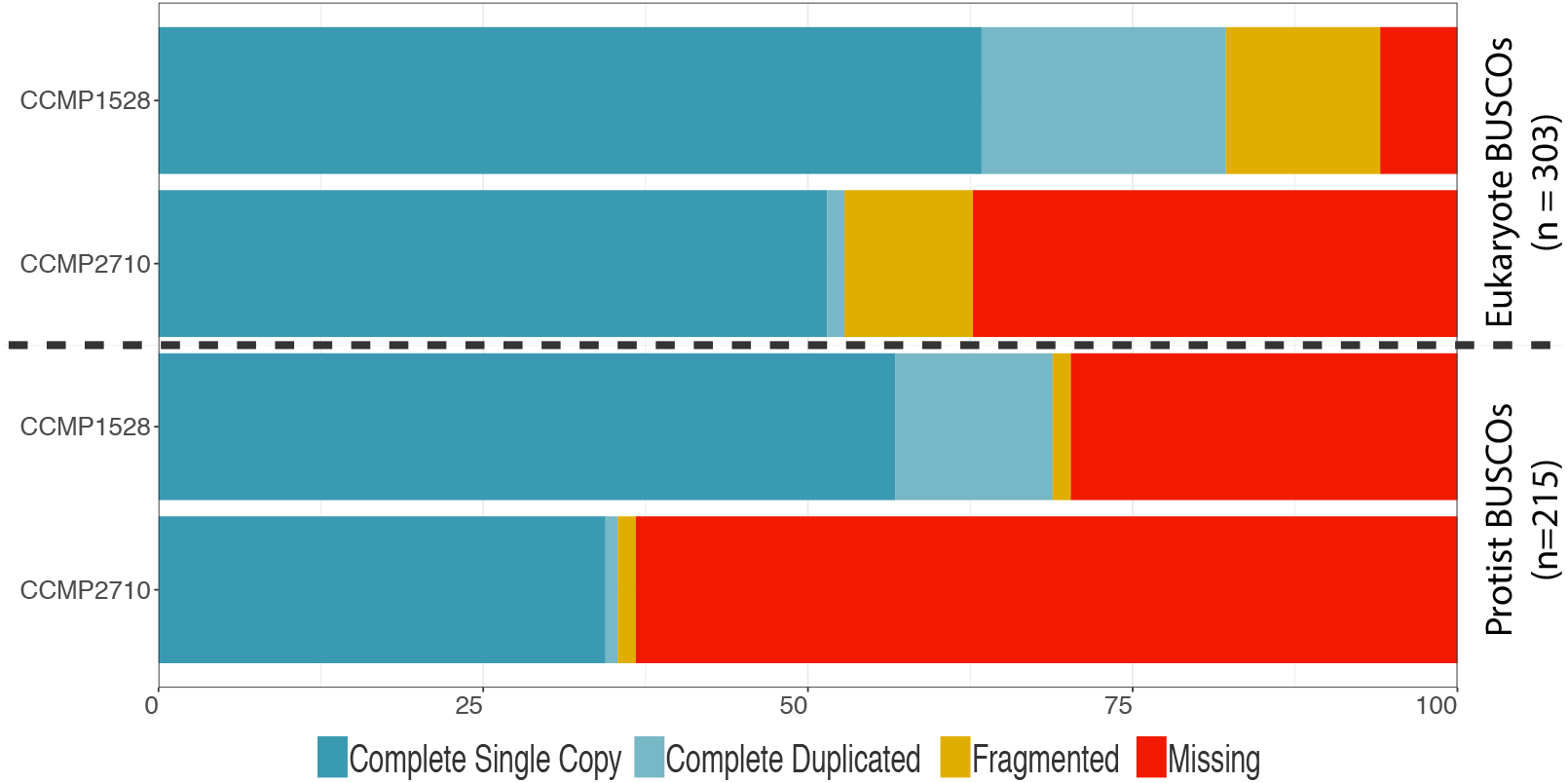
Percent of eukaryote and protist Benchmarking Universal Single Copy Orthologs (BUSCOs) complete, fragmented, or missing in *Phaeocystis globosa* CCMP1528 and *Phaeocystis sp.* CCMP2710 transcriptomes. BUSCO software was used to determine the percent of eukaryotic and protistan BUSCOs represented by complete single copies, complete but duplicated copies, copies that were fragmented or missing in the *Phaeocystis globosa* CCMP1528 transcriptome assembled for this study and the *Phaeocystis sp.* CCMP2710 transcriptome assembled for the Marine Microbial Eukaryote Transcriptome Sequencing Project (MMETSP). More eukaryotic and protistan BUSCOs were represented in the *Phaeocystis globosa* CCMP1528 transcriptome than the *Phaeocystis sp.* CCMP2710 transcriptome. The plot was rendered with the R package ggplot2.

Annotation was possible for relatively few of the contigs in the *P. globosa* CCMP1528 transcriptome assembly, but more genes were annotated with the Pfam and GO annotation pipeline (26%) than with the KEGG pipeline (14%). Additionally, both annotation methods annotated the significantly differentially expressed (DE) genes at a higher rate than the whole transcriptome (Table 1).

**Table 1.** Pfam, Gene Ontology (GO), and Kyoto Encyclopedia of Genes and Genomes (KEGG) annotation statistics for the *Phaeocystis globosa* CCMP1528 transcriptome assembly and differentially expressed (DE) genes.

|  | Pfam / GO | KEGG |
| --- | --- | --- |
| Total genes annotated | 17,826 (26%) | 9,967 (14%) |
| Total w. pathway or GO | 8,962 (13%) | 5,585 (8%) |
| DE genes annotated | 5,180 (66%) | 3,305 (43%) |
| DE genes w. pathway or GO | 2,764 (35%) | 1,883 (24%) |

### Differential gene expression analysis

Gene expression patterns in colonial and solitary replicates were explored with a principal component analysis (PCA) and an expression heatmap. Initial data exploration revealed the colonial replicate ‘C2’ as an outlier to other colonial replicates and solitary replicates (Fig. S5A, B) and this sample was excluded from further analyses. The remaining 3 colonial replicates clustered separately from the 4 solitary replicates in a PCA plot (Fig. 2A). Differential expression analysis identified 535 genes as significantly upregulated and 7,234 genes as significantly downregulated in colonial replicates. An expression heatmap of the most differentially expressed genes sorted by FDR adjusted p-value (padj) clearly illustrates the overall expression pattern—the majority of significantly differentially expressed genes are downregulated in colonial replicates (Fig. 2B).

**Figure 2.**
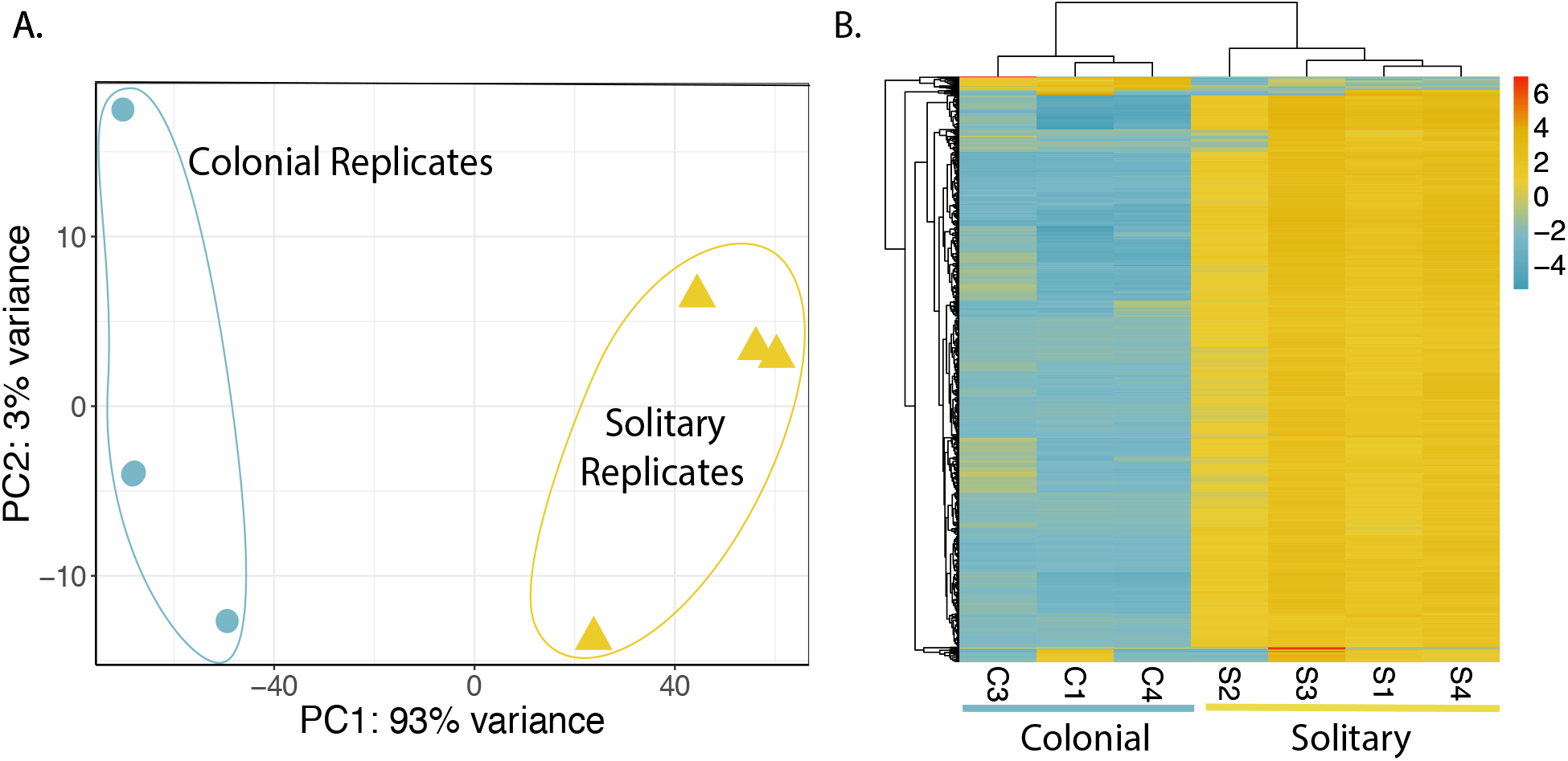
Principal component analysis (PCA) and heatmap demonstrating gene expression patterns in colonial and solitary *Phaeocystis globosa*. **A.** PCA performed on distances between samples derived from regularized log transformed counts. Colonial and solitary replicates cluster separately, and the majority of variance is between sample type rather than within replicates. Results plotted with R package ggplot2. **B.** Heatmap includes the 1000 significantly differentially expressed genes with the lowest FDR adjusted p-values. Heatmap color represents difference from the mean regularized log transformed count for each contig in each sample. The majority of differentially expressed genes are downregulated in colonial replicates, and replicates cluster by sample type. Results plotted with R package pheatmap.

### Gene set enrichment analysis

In order to identify Biological Process (BP) GO terms over-represented in significantly up- and downregulated gene sets, we applied a hypergeometric test with a significance cut-off of p < 0.05. Twenty BP GO term were over-represented among significantly upregulated genes and were primarily involved in cell signal transduction in response to external stimuli (Fig. 3; Table S3). Notably, GO terms involving arabinose, a component of the colonial matrix, were also enriched among upregulated genes. In the downregulated gene set, 48 BP GO terms were enriched, including several involved in cation transport, response to oxygen-containing compounds, translation and protein transport, and vacuolar transport and exocytosis (Fig. 3; Table S4). REVIGO software was used to remove redundant GO terms from lists of enriched terms and to visualize results in a Multidimensional Scaling (MDS) plot (Fig. 3) based on GO term semantic similarities (Supek et al. 2011) as determined by shared ancestry (Pesquita et al. 2009).

**Figure 3.**
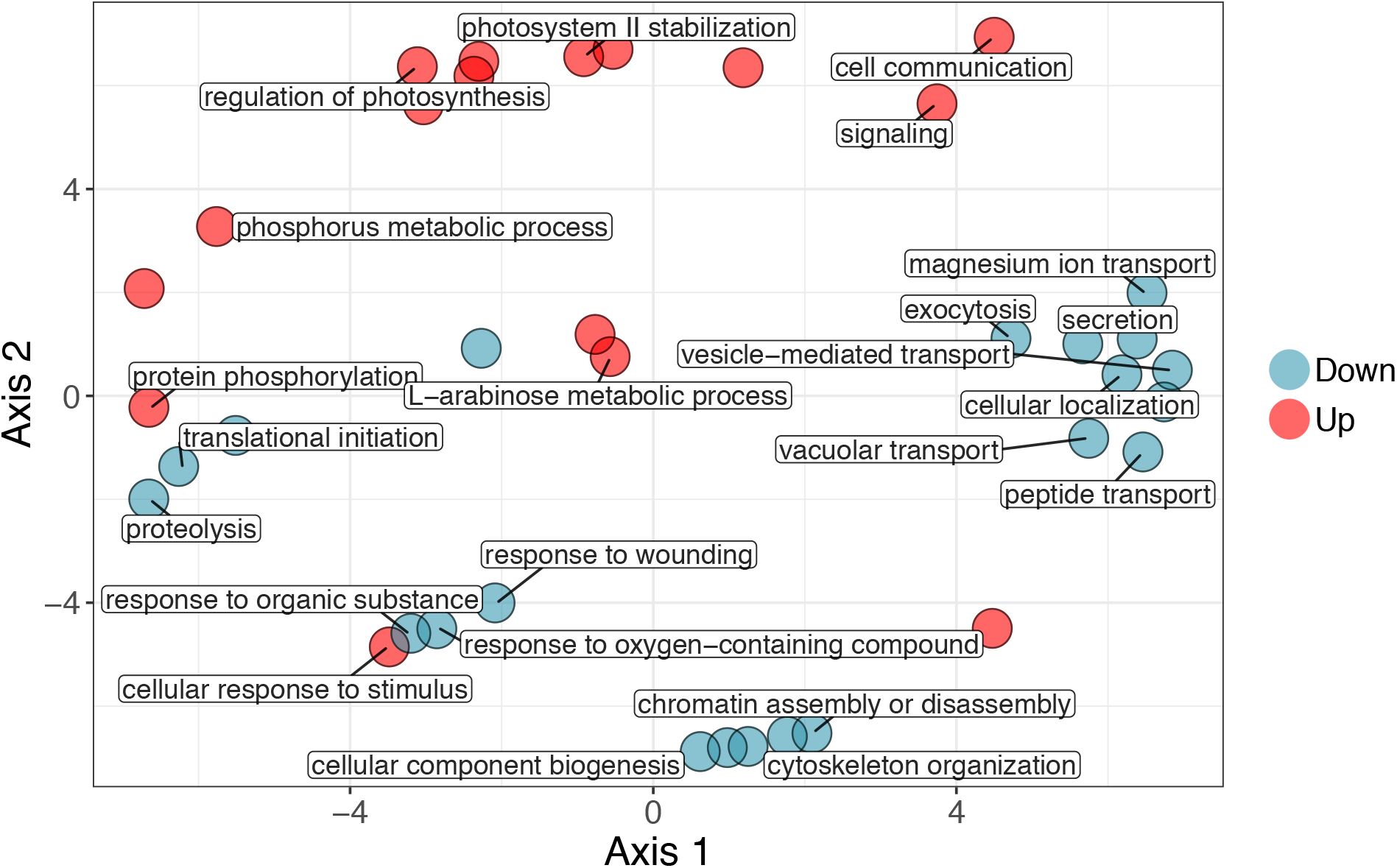
Multidimensional scaling plot of semantic similarities between non-redundant GO terms over-represented in significantly up- and downregulated gene sets. Analysis performed using the REVIGO tool (http://revigo.irb.hr/) with the allowed similarity set to 0.7 (to remove redundant GO terms) and the SimRel metric selected to calculate similarities. REVIGO results were exported to the R statistical environment and plotted with ggplot2. Representative GO terms were manually selected and labeled on the plot. To view all GO term labels, an interactive version of the plot made with R package ggplotly is available at: https://brisbin.shinyapps.io/shinycolsol/.

When we applied hypergeometric testing to KEGG pathways, only the cGMP-PKG signaling pathway (cyclic guanosine monophosphate-protein kinase G pathway) was enriched in the upregulated gene set (p<0.05) (Table S5). Five pathways were enriched among downregulated genes: Lysosome, Autophagy, MAPK signaling pathway (mitogen activated protein kinase signaling pathway), AMPK signaling pathway (adenosine monophosphate-activated protein kinase signaling pathway), and Epidermal growth factor receptor (EGFR) tyrosine kinase inhibitor resistance (Table S6). The lower annotation rate for KEGG pathways compared with GO terms contributed to the difference in enrichment testing results. We therefore also applied a linear model test for KEGG pathway enrichment, which identified several additional pathways as being significantly enriched in the up- (6) and downregulated (7) gene sets. With this additional test, the PI3K-Akt signaling (phosphoinositide 3-kinase-protein kinase B signaling pathway), Glycosphingolipid biosynthesis, Ferroptosis, Plant-pathogen interaction, Circadian rhythm, Viral carcinogenesis pathways were also enriched among upregulated genes (Table S7). The Protein processing in endoplasmic reticulum, Oxidative phosphorylation, Ras signaling pathway, Sphingolipid metabolism, Steroid biosynthesis, Fatty acid degradation, and Taste transduction pathways were additionally identified as enriched in downregulated genes (Table S8).

### Genes associated with DMSP and DMS production

A blastp query against curated DSYB protein sequences from Curson et al. (2018) and Alma family protein sequences from Alcolombri et al. (2015) identified 4 *Phaeocystis globosa* contigs as putative *DSYB* or *Alma* family genes (Table 2). The *P. globosa* amino acid (AA) sequence translated from Transcript_30752 aligned with the sequence for *Prymnesium parvum* CCAP946/1B DSYB protein, which is experimentally proven to be highly active. It is therefore likely that this gene is actively involved in DMSP biosynthesis in *Phaeocystis globosa*. A second *P. globosa* AA sequence, from Transcript_31221, aligned with the *Pseudonitzchia fraudulenta* DYSB protein sequence, making it also a possible *DSYB* gene. The *Pseudonitzchia* DSYB protein has not been experimentally proven to be active, but its sequence is phylogenetically close to the *Fragillariopsis* DSYB, which has been proven to be active. Neither putative *P. globosa DSYB* genes were differentially expressed between solitary and colonial culture replicates in this study, but both were expressed at relatively high levels in both sample types (Fig. 4).

**Table 2.** Top *Phaeocystis globosa* CCMP1528 blastp results against DSYB and Alma family reference sequences (e-values <1E-30).

|  | Database Sequence | %ID | Alignment Length (AAs) | Gaps | E-value | Bit Score |
| --- | --- | --- | --- | --- | --- | --- |
| Transcript_30752 | *Prymnesium parvum* CCAP946/1B DSYB | 74 | 314 | 2 | 1.15E-170 | 472 |
| Transcript_31221 | *Pseudonitzschia fraudulenta* WWA7 DSYB | 45 | 263 | 8 | 1.92E-65 | 210 |
| Transcript_36000 | *Emiliana huxleyi* Alma7 | 61 | 272 | 4 | 3.70E-119 | 336 |
| Transcript_68879 | *Emiliana huxleyi* Alma4 | 40 | 253 | 4 | 2.64E-48 | 166 |

**Figure 4.**
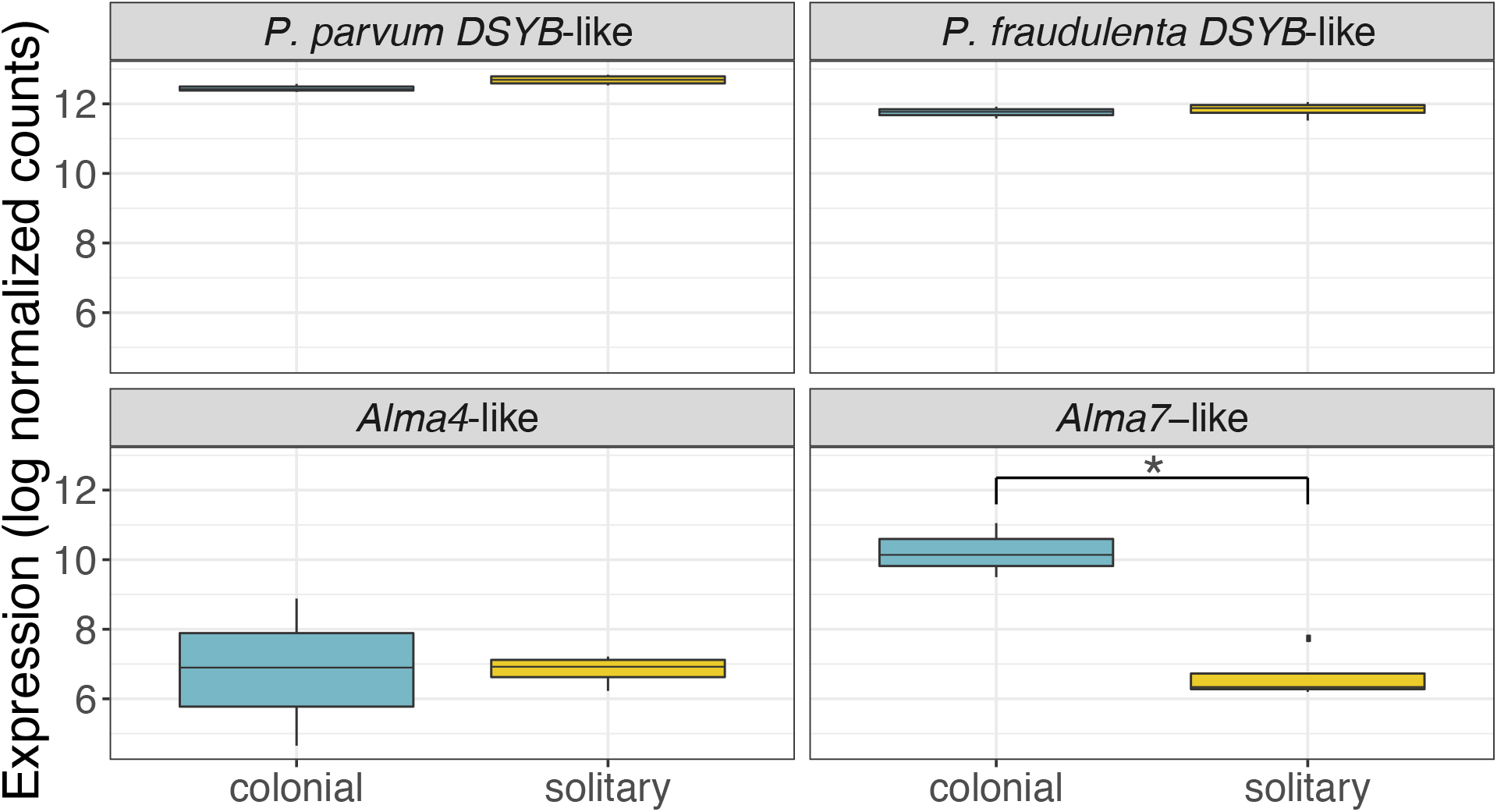
Normalized expression levels of *DSYB*-like and *Alma*-like genes in *Phaeocystis globosa* colonial and solitary cell cultures. Box plots show the range, quartiles and median of the log normalized counts for each gene in colonial and solitary culture replicates. Only the *E. huxleyi Alma7*-like gene was significantly differentially expressed and was upregulated in colonial samples (padj = 1.44E-12, log fold-change = 3.55) and is represented with an asterisk (*) on the plot. Plots made with R package ggplot2.

Two *Alma* family-like genes were identified in the *Phaeocystis globosa* transcriptome. One *P. globosa* AA sequence, from Transcript_36000, aligned with *Emiliania huxleyi* Alma7. All 4 *Alma* homologs identified from the MMETSP *Phaeocystis antarctica* transcriptome are phylogenetically closest to the *E. huxleyi Alma7*, but *E. huxleyi* Alma7 has not been proven to have DMSP-lyase activity. Another *P. globosa* AA sequence, from Transcript_68879, aligned with *E. huxleyi* Alma4, which also has not been experimentally proven active. Both putative *P. globosa Alma* family genes were expressed at lower rates than *DSYB*-like genes. The *P. globosa Alma4*-like gene was not differentially expressed in solitary and colonial culture replicates in this study (Fig. 4). The *P. globosa Alma7*-like gene, however, was significantly upregulated in colonial replicates (padj = 1.44E-12, logFC = 3.55), suggesting that DMSP biosynthesis is occurring in both colonial and solitary cells, but colonial cells may be cleaving DMSP to DMS and acrylate more actively than solitary cells.

## DISCUSSION

Colonial *Phaeocystis* blooms widely impact ecosystem function and can be extremely detrimental in some systems, particularly in subtropical and tropical coastal regions. Although many hypotheses exist, factors initiating colonial blooms and the ecological function of colonial formation remain enigmatic. We investigated gene expression associated with colony formation in a warm-water colony-forming strain of *Phaeocystis globosa* to identify cellular processes associated with colony formation and potentially provide clues for what initiates colony formation and the functional role of colonies in the *Phaeocystis* life-cycle. Overall, we observed a transcriptional shift in colonial cultures compared to solitary cell cultures, with vastly more genes significantly downregulated in colonial cells than upregulated (Fig. 2). This shift suggests that there are trade-offs associated with colony production and resources must be diverted to construct and maintain the colonial matrix. A relatively small number of genes are upregulated to produce colonies, but the low annotation rate of these genes, and the transcriptome overall, make it challenging to fully interpret the results (Table 1). Gene set enrichment analyses inherently relies on how many and which genes are annotated and systematic biases likely in gene annotation will influence results (Haynes et al. 2018). However, these analyses still assist in identifying pathways and functions that may be important to the question at hand and indicate genes and pathways that should be followed up in future studies. The results presented here highlight genes involved in constructing the colonial matrix, changes in cellular morphology, responding to external stimuli, cellular proliferation, and producing DMSP, DMS, and acrylate. Results from this study support a defensive role for colony formation in *Phaeocystis globosa*.

### Colony matrix carbohydrates and colonial cell morphology

Differential expression of genes associated with clearly observable changes between treatment groups can serve to “ground-truth” results from RNA-seq experiments and therefore increase confidence in expression changes detected for genes for less observable traits. In this study, changes in cellular morphology and colony formation itself are clearly observable differences for which several associated genes are differentially expressed. The observed expression patterns for these genes can additionally provide new insight into the construction of the colonial matrix and pathways associated with morphological changes in colonial *Phaeocystis* cells. Many different polysaccharides are recognized as contributors to the matrix structure of *Phaeocystis globosa* colonies, including arabinose, rhamnose, xylose, mannose, galactose, glucose, gluconuronate, and O-methylated pentose sugars (Janse et al. 1996). *Phaeocystis* isolated from different locations tends to have distinct matrix carbohydrate fingerprints, which may be due to genetic attributes of different strains or which could arise from different environmental conditions, such as light or nutrient availability. For example, arabinose is the most abundant matric carbohydrate in *P. globosa* sampled from the North Sea (Janse et al. 1996). In this study, the GO term for Arabinose metabolic process was enriched among upregulated genes in colonial cells (Fig. 3; Table S3). These results indicate that arabinose is likely the dominant matrix polysaccharide in *P. globosa* CCMP1528 and that arabinose production is specifically associated with colony formation in this strain. The colonial matrix also contains nitrogen (Hamm 2000), which is likely included in amino sugars (Solomon et al. 2003). Our results, however, did not indicate that amino sugar biosynthesis or metabolism was upregulated in colonial cells.

Divalent cations, particularly Mg^2+^ or Ca^2+^, are required for colonial polymers to gel and contribute to the stability of the colonial matrix (van Boekel 1992). GO terms for Divalent inorganic cation transport, Magnesium ion transport, and Divalent metal ion transport, however were enriched in downregulated genes in colonial replicates (Fig. 3; Table S4). Similarly, Bender et al. (2018) found that *Phaeocystis antarctica* produces more calcium-binding proteins when iron limitation decreases colony formation. These results may be due to the importance of divalent cations for flagellate motility. Actively swimming *Phaeocystis globosa* flagellates, as observed in this study, may require continuous transport of divalent cations to the point of masking their shared importance in colony formation. Calcium signaling also induces secretion of vesicles containing gel forming polymers (Chin et al. 2004), which further confounds the observed downregulation of divalent cation transport genes in colonies. However, flagellates also secrete vesicles, but instead of gel polymers they contain star-shaped structures composed of chitinous filaments (Chretiennot-Dinet et al. 1997). The exact function of these structures is unknown, but they may be involved in mating or defense (Dutz & Koski 2006). In addition to divalent cation transport, a number of other GO-terms enriched in downregulated genes may be involved in secreting these structures, such as Exocytosis, Secretion, Vesicle mediated transport, and Vacuolar transport (Fig. 3; Table S3). Alternatively, these GO terms may be involved in scale formation and secretion (Taylor et al. 2007), as scales are only observed on *Phaeocystis globosa* flagellates and not colonial cells (Rousseau et al. 2007).

### A defensive role for colony formation: resource allocation, pathogen interaction, and DMS/acrylate production

Out of 7,769 genes that were significantly differentially expressed between colonial and solitary replicates, 7,234 genes were downregulated in colonial cells. This dramatic transcriptional shift in colonial cells supports a high resource cost associated with producing colonies (Wang et al. 2015). Specifically, our results indicate that resources are being diverted from protein translation and transport and cell division in order to produce the colonial matrix. Several GO terms involved in the synthesis of larger nitrogenous compounds and their transport, including Translation initiation, Protein metabolic process, Protein N-linked glycosylation, and Protein transport were significantly enriched in downregulated genes in colonial cells (Fig. 3; Table S4). Similarly, the KEGG pathway, Protein processing in endoplasmic reticulum, was also enriched in downregulated genes in colonies (Table S7). Likewise, several mitosis-associated GO terms (Chromatin assembly and disassembly, Cytoskeleton organization, Cellular component biogenesis) were also enriched among downregulated genes in colonial cells (Fig. 3; Table S4). However, the downregulation of mitosis-associated genes in colonial cells conflicts with observations in previous studies. Veldhius et al. (2005) observed that colonial cells divide at a higher rate than solitary cells and proposed that in addition to experiencing less grazing and viral lysis, colonial cells may dominate blooms because they outgrow solitary cells. We believe the difference in our results may be due to the type of solitary cells observed—the solitary cells in previous studies could have been diploid flagellates, especially if they were derived from disrupted colonies, whereas solitary cells in our study are likely haploid flagellates, which have been reported to divide extremely rapidly (Rousseau et al. 2007).

There were also several signaling pathways represented in the results suggesting that colonial cells are exposed to fewer general stressors, but may be responding to more strongly to specific pathogens. In plants, the MAPK (mitogen activated protein kinase) pathway primarily transduces signals from extracellular stressors to the nucleus or cytoplasm and initiates an appropriate response (Taj et al. 2010). In our results, the MAPK signaling pathway was significantly enriched in genes downregulated in colonies. The downregulated genes in this pathway encode MAP3Ks, MAP2Ks, and MAPKs, which are activated in response to pathogen attack and infection, phytohormones, cold and salt stress, and reactive oxygen species (Taj et al. 2010). Downregulation of genes associated with stress response is also evidenced by related GO terms enriched among downregulated genes, specifically Response to wounding and Response to oxygen-containing compounds. These results support the hypothesis that colony formation serves a defensive purpose. Defense responses regulated by the MAPK pathway are unneeded because the colony skin is protecting cells from these stressors. However, the Plant-pathogen interaction pathway was enriched in upregulated genes in colonial cells, indicating that specific pathogens may penetrate the colonial fortress or that pathogen interaction may play a role in stimulating colony formation. Genes upregulated in this pathway were for calcium-dependent protein kinases and calcium-binding protein CML (calmodulin-like protein), immune response genes that are activated following recognition of specific pathogen-associated molecular patterns (Cheval et al. 2013). Specific bacterial interactions are known to influence transitions between life-cycle stages in several other protists: specific bacteria stimulate growth in marine diatoms (Amin et al. 2015) and specific bacterial signaling molecules are responsible for inducing both colony formation (Woznica et al. 2016) and sexual reproduction (Woznica et al. 2017) in choanoflagellates, another single-celled colony-forming marine plankton. In *Phaeocystis*, axenic cultures exhibit decreased growth rates (Solomon et al. 2003), but the effects of specific bacteria on colony formation have not yet been investigated.

*Phaeocystis* is a copious producer of DMSP and its cleavage products, DMS and acrylate. DMS and acrylate have been indicated as grazer-deterrents and antimicrobials (Hamm 2000; Noordkamp et al. 2000; Wolfe & Steinke 1996). While algal genes associated with DMSP biosynthesis (*DSYB*, Curson et al. 2018) and cleavage (*Alma* family genes, Alcolombri et al. 2015) have been identified in many algal transcriptomes, including *Phaeocystis antarctica*, this study is the first to identify these genes for *Phaeocystis globosa*. We found that colonial and solitary *P. globosa* expressed *DSYB*-like genes at the similar levels, suggesting that the two cell types produce similar amounts of DMSP. *DSYB* expression in *Prymnesium parvum*, the haptophyte in which DSYB was discovered, is affected only by salinity, potentially indicating that DMSP production functions primarily in osmoregulation rather than as a defensive or stress response (Curson et al. 2018). Similar *DSYB*-like gene expression levels in colonial and solitary *P. globosa* cells support a basic, shared function for DMSP in the two cell types. Contrastingly, an *Alma* family gene was upregulated in colonial cells. Acrylate accumulates in *Phaeocystis* colonies and may serve to deter grazers and pathogens from disrupting the colonial matrix (Noordkamp et al. 2000). While acrylate may accumulate in colonies simply because it cannot escape through the colonial skin, the upregulation of an *Alma*-like gene in colonies suggests that colonial *Phaeocystis* cells may actively produce excess acrylate and DMS than solitary cells, further supporting a defensive role for colony production in *P. globosa*.

### Role of colonies in *Phaeocystis* reproduction

Colony formation is believed to be involved in sexual reproduction in *Phaeocystis* since swarming flagellates have been observed within senescent colonies (Peperzak et al. 2000; Rousseau et al. 2013). However, we did not find meiosis or sexual reproduction GO terms or KEGG pathways enriched in up- (or down-) regulated genes in this study. These results may arise from RNA being extracted during mid- to late exponential growth phase. Previous observations suggest that colonies produce flagellates during bloom decay, so we might have found meiosis genes upregulated in colonial cells if we had sampled toward late stationary phase instead of exponential phase.

### Signaling pathways associated with colony formation

Processes and pathways involved in cell-signaling, cell communication, and response to stimuli that are enriched in upregulated-genes are particularly interesting because they shed some light on factors stimulating colony formation in *Phaeocystis globosa*. The cGMP-PKG signaling pathway was the only KEGG pathway significantly enriched among upregulated genes when a hypergeometric test was used. Three genes in this pathway were upregulated: 1) cGMP-dependent protein kinase, which phosphorylates biologically important targets, has been implicated in cell division and nucleic acid synthesis, and reduces cytoplasmic Ca^2+^ concentrations (Lincoln et al. 2001); 2) cAMP-dependent protein kinase regulator; and 3) a cAMP-responsive element-binding protein (CREB), which binds to DNA to increase or decrease transcription and is associated with increased cell survival (Chrivia et al. 1993). The PI3K-Akt signaling pathway was also significantly enriched in upregulated genes when the additional linear model test was used. Within this pathway, two Extracellular Matrix (ECM) focal adhesion genes, for Tenascin (a glycoprotein) and Type IV collagen, were significantly upregulated. Focal adhesion proteins connect cells to extracellular matrices both literally and figuratively, by holding cells in place and by initiating cellular responses to external conditions (Wozniak et al. 2004). Bender et al. (2018) also found focal adhesion proteins, specifically glycoproteins, upregulated in colonial *Phaeocystis antarctica*. It is therefore likely that these proteins have an important function in structurally maintaining cell positions in the colonial matrix and signaling between colonial cells. Focal adhesion proteins may be mediating interactions with protein kinases in colonial cells, which go on to promote cell proliferation and differentiation into the colonial morphotype. These signaling pathways represent important candidates for continued study of molecular mechanisms regulating colony formation.

### Conclusions and future directions

This study investigated gene expression associated with colony formation in *Phaeocystis globosa* for the first time and discovered a large transcriptional shift associated with colony production. Differentially expressed genes were mostly downregulated in colonies, providing evidence for extensive resource allocation toward colony formation. Together, activation of pathogen interaction pathways, reduced expression of stress-response pathways, and increased expression of a DMSP-lyase, which produces DMS and acrylate, supporting a defensive role for colony formation. Future studies may extend this work by investigating *P. globosa* gene expression in colonial and solitary cells in a time course study through the waxing and waning of a bloom and under different nutrient and grazing regimes, potentially by using mesocosms or metatranscriptomic methods in natural communities. While our ability to fully interpret the results was inhibited by an overall lack of annotated genomes and transcriptomes for diverse protist lineages, this study represents a step in the right direction by contributing a new and deeply sequenced transcriptome for *Phaeocystis globosa*. Identification of *DSYB* and *Alma* family-like genes in this transcriptome will additionally allow for further investigation into *P. globosa* DMSP and DMS production in the oceans. We were also able to identify several protein kinase signaling pathways that are potentially important for regulating colony formation and should be experimentally investigated in follow-up studies. The results presented here will guide and facilitate continued efforts to unravel the complex factors responsible for triggering harmful colonial *Phaeocystis* blooms, which will likely increase with continued climate change and nutrient pollution in the future.

## ACKNOWLEDGEMENTS

This work was supported by funding from the Marine Biophysics Unit of the Okinawa Institute of Science and Technology Graduate University. MMB is supported by a JSPS DC1 graduate student fellowship. We thank Angela Ares Pita for valuable discussion and suggestions and for helpful feedback on manuscript drafts. Hiroki Goto and the OIST sequencing section provided important guidance for RNA library preparation and other sequencing considerations. Steven D. Aird edited the manuscript and provided helpful comments.

## DATA AVAILABILITY

Sequence data is available in the NCBI Sequence Read Archive (SRA) with accession numbers SRR7811979–SRR7811986. The *Phaeocystis globosa* CCMP1528 *de novo* transcriptome produced and used in this study, data files, and data analysis scripts can be accessed at https://doi.org/10.5281/zenodo.1476491.

